# High-throughput amino acid-level characterization of the interactions of plasminogen activator inhibitor-1 with variably divergent proteases

**DOI:** 10.1101/2024.09.16.612699

**Authors:** Laura M Haynes, Matthew L Holding, Hannah DiGiovanni, David Siemieniak, David Ginsburg

## Abstract

While members of large paralogous protein families share structural features, their functional niches often diverge significantly. Serine protease inhibitors (SERPINs), whose members typically function as covalent inhibitors of serine proteases, are one such family. Plasminogen activator inhibitor-1 (PAI-1) is a prototypic SERPIN, which canonically inhibits tissue-and urokinase-type plasminogen activators (tPA and uPA) to regulate fibrinolysis. PAI-1 has been shown to also inhibit other serine proteases, including coagulation factor XIIa (FXIIa) and transmembrane serine protease 2 (TMPRSS2). The structural determinants of PAI-1 inhibitory function toward these non-canonical protease targets, and the biological significance of these functions, are unknown. We applied deep mutational scanning (DMS) to assess the effects of ∼80% of all possible single amino acid substitutions in PAI-1 on its ability to inhibit three putative serine protease targets (uPA, FXIIa, and TMPRSS2). Selection with each target protease generated a unique PAI-1 mutational landscape, with the determinants of protease specificity distributed throughout PAI-1’s primary sequence. Next, we conducted a comparative analysis of extant orthologous sequences, demonstrating that key residues modulating PAI-1 inhibition of uPA and FXIIa, but not TMPRSS2, are maintained by purifying selection. PAI-1’s activity toward FXIIa may reflect how protease evolutionary relationships predict SERPIN functional divergence, which we support via a cophylogenetic analysis of all secreted SERPINs and their cognate serine proteases. This work provides insight into the functional diversification of SERPINs and lays the framework for extending these studies to other proteases and their regulators.

## Introduction

Large paralogous protein families in which members share a common protein structure are ubiquitous across all clades of life. Diversity often results from gene duplication events that occurred during either whole genome duplication or as segmental duplications of one or more genes (Hahn 2009). To acquire and maintain distinct functions despite similarities in sequence and structure, members of these paralogous protein families must acquire unique functional niches (McClune, et al. 2019; McClune and Laub 2020; Nocedal and Laub 2022). Revealing the shape and complexity of these niches is crucial to understanding the emergence and insulation of protein-protein interactions.

**<u>Ser</u>**ine **<u>p</u>**rotease **<u>in</u>**hibitors, or SERPINs, are a protein superfamily with high functional variability and specificity. Inhibitory SERPINs are defined by their ability to efficiently and specifically inhibit one or more members of the highly paralogous serine protease protein family. Both protein families are found across all domains of life, where SERPINs covalently bind serine proteases, irreversibly inhibiting them (Spence, et al. 2021). In humans and other vertebrates, SERPINs have been shown to regulate a range of processes, including immune/inflammatory responses, coagulation/fibrinolysis, and extracellular matrix remodeling (Sanrattana, et al. 2019). While all SERPINs share a common protein fold, most SERPINs perform their biological function as specific and irreversible inhibitors of serine proteases via a “mousetrap” mechanism (**Fig. 1**) (Law, et al. 2006).

**Figure 1.**
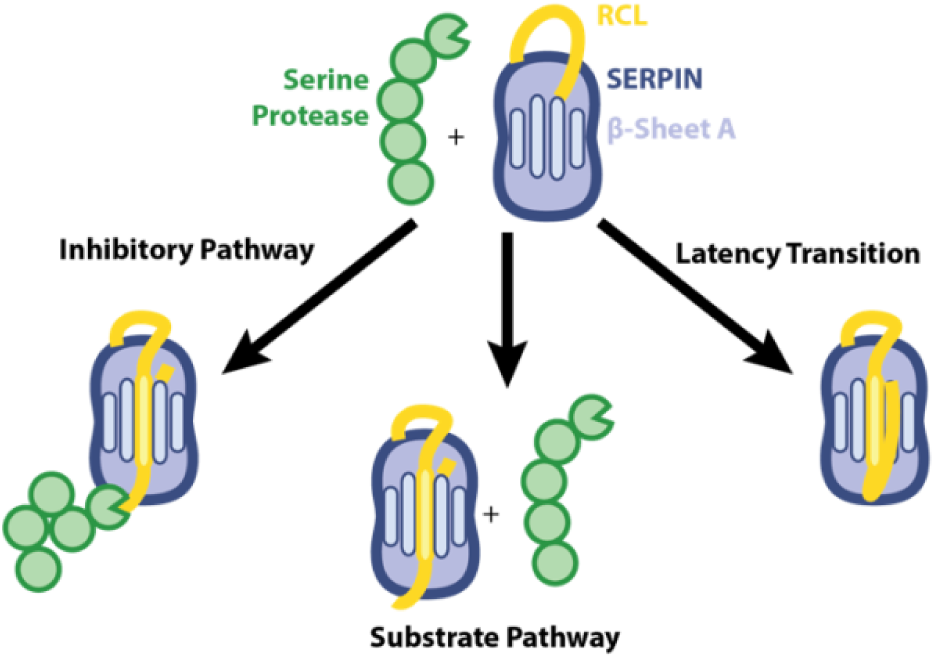
SERPINs inhibit serine proteases via a “mousetrap” mechanism to inhibit their primary and non-canonical protease targets. In its metastable active state, the SERPIN’s reactive center loop (RCL, yellow) extends from its central globular core and contains a preferred cleavage site for proteases targeted for inhibition (green). Under conditions favoring the inhibitory pathway (*left pathway*), after the protease cleaves the scissile bond in the RCL, the acyl intermediate is trapped when the RCL inserts into β-sheet A forming a fifth β-strand. Alternatively, the protease cleaves the RCL and the acyl intermediate is hydrolyzed resulting in the insertion of the RCL into β-sheet A and a non-functional SERPIN, while the protease remains active (*center pathway*). Unique among SERPINs, PAI-1 can also undergo a spontaneous latency transition in which the RCL inserts into β-sheet in the absence of a proteolytic event, rendering the SERPIN inactive (*right pathway*).

In their active state, SERPINs exist in a metastable conformation in which the reactive center loop (RCL), which contains an amino acid sequence that mimics that of their target serine protease’s preferred cleavage site, extends from the globular core of the protein. Upon SERPIN/protease engagement, irreversible inhibition of the serine protease occurs when the RCL inserts into the SERPIN’s central β-sheet A prior to the resolution of the acyl intermediate; however, if hydrolysis of the acyl intermediate occurs prior to RCL insertion, the SERPIN serves as a substrate rather than an inhibitor of the serine protease–both of these conformations occupy a lower energy space (**Fig. 1**) (Lawrence, et al. 1995; Gettins 2002; Law, et al. 2006; Huntington 2011). Although a SERPIN’s protease specificity is derived in part from the RCL amino acid sequence, structural features located distal to the RCL also contribute significantly (Gettins and Olson 2009; Marijanovic, et al. 2019). Some SERPINs appear to be highly specific inhibitors of a single protease, while others exhibit activity toward additional protease targets, often with decreased efficiency compared to their canonical protease target (Gettins 2002). Exosite interactions, as well as a limited number of mutations, are also known to alter SERPIN inhibitory profiles (Gettins 2002; Huntington 2011). The specificity of SERPINs for their target serine protease(s) therefore provides not only a novel system for studying the development of functional niches in the context of paralog evolution, but also one where the determinants of success or failure of the protein-protein interaction are complex.

Plasminogen activator inhibitor-1 (PAI-1, encoded by the *SERPINE1* gene) is a 379 amino acid SERPIN canonically considered to be a regulator of fibrinolysis, the enzymatic degradation of fibrin clots, by inhibiting two closely related proteases, urokinase and tissue-type plasminogen activators (uPA (encoded by the *PLAU* gene) and tPA (encoded by the *PLAT* gene), respectively) with high efficiency (second-order rate constants of ∼10^6^-10^7^ M^-1^ s^-1^, (Sherman, et al. 1992)) and a stoichiometry of inhibition (SI) of approximately one (Lawrence, et al. 2000). PAI-1 has also been identified as a possible inhibitor of other hemostatic proteases, including the procoagulant factor (F) XIIa (encoded by the *F12* gene), which stands in stark juxtaposition to PAI-1’s primary function as an inhibitor of the clot dissolving proteases tPA and uPA (Berrettini, et al. 1989; Keijer, et al. 1991; Rezaie 2001; Tanaka, et al. 2009; Sen, et al. 2011; Puy, et al. 2019). PAI-1 also inhibits transmembrane serine protease 2 (TMPRSS2, **SI Fig. 1**), through which it mediates the course of viral infections such as influenza and SARS-CoV-2 (Dittmann, et al. 2015; Shen, et al. 2017), with increased PAI-1 levels reported in response to influenza (Keller, et al. 2006; Bouwman, et al. 2009) and SARS-CoV-2 infections (Al-Samkari, et al. 2020; Zuo, et al. 2021).

In the present work, we use deep mutational scanning (DMS) to analyze the mutational landscape of PAI-1 inhibition of several proteases, including uPA and two non-canonical targets of PAI-1 (FXIIa and soluble TMPRSS2), characterizing the overall degree and potential regionalization of marginal specificity in these interactions. We follow prior work that defines protein sequence space as the set of all possible amino acids in a protein coupled with the possible evolutionary trajectories to a new sequence (McClune and Laub 2020). Within a sequence space, *robust* and *marginal specificity* refer to members of paralogous protein superfamilies for which multiple mutations are required to alter their specificity and cases where a single amino acid substitution can alter a paralog’s specificity, respectively (Ghose, et al. 2023). We then use comparative evolutionary analyses to contextualize the results of our DMS screen to explore the relationships between protease conservation, paralogous protein relationships, and the sequence space in which these proteins interact.

## Results and Discussion

### Construction and characterization of a PAI-1 variant library on the I91L background

We constructed a novel phage display library as previously described (Huttinger, et al. 2021; Haynes, et al. 2022), though with the addition of an I91L PAI-1 variant backbone to increase the half-life of PAI-1 in its metastable active conformation from 1-2 h to ∼19 h (**Fig. 1**) (Berkenpas, et al. 1995; Haynes, et al. 2022). The I91L background was chosen to isolate the effects of mutants on PAI-1 serine protease target specificity from those impacting functional stability.

The I91L PAI-1 variant library exhibited a depth of 6.6x10^6^ unique phage clones with an average of three missense or nonsense mutations per clone (**SI Fig. 2A**). Of the 7,201 possible single amino acid substitutions in PAI-1 (379x19), 5,688 (79%) are represented in the I91L variant library as determined by high-throughput sequencing (HTS). The abundance of each variant is highly correlated (slope = 0.95, R^2^ = 0.85; **SI Fig. 2B**) with our previous variant library on the wild-type (WT) PAI-1 background (Huttinger, et al. 2021). Log_2_-fold enrichment scores for uPA functional inhibitory variants in the I91L PAI-1 library are also highly correlated with those previously determined (Huttinger, et al. 2021) on the WT PAI-1 background (**SI Fig. 2C**; R^2^ = 0.66, slope = 0.45).

**Figure 2.**
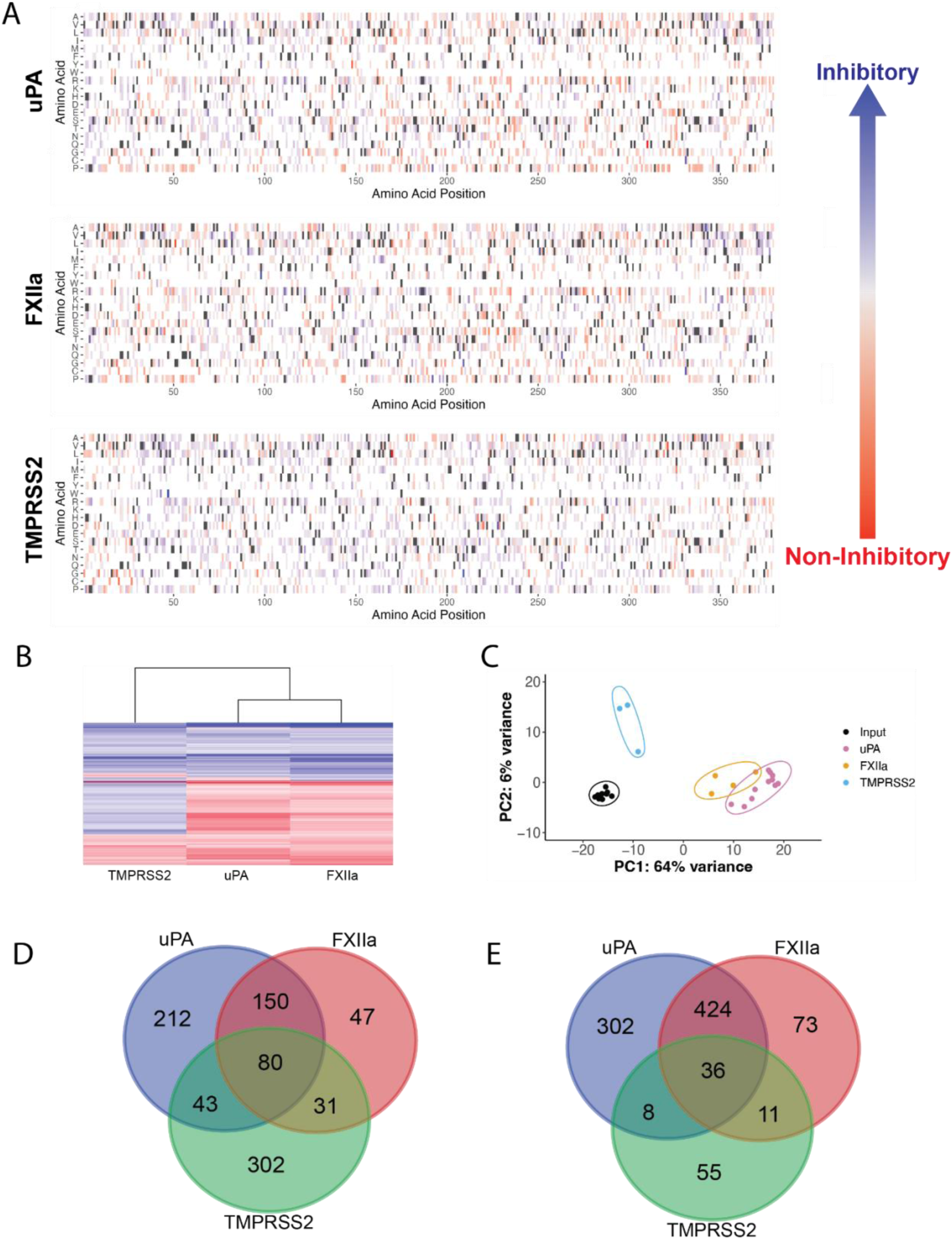
PAI-1 DMS and differential patterns of serine protease inhibition. (A) PAI-1 specificity fingerprints are shown for uPA, FXIIa, and TMPRSS2. The PAI-1 amino acid sequence is shown along the x-axis with potential amino acid substitutions shown along the y-axis. Tolerated mutations are shown in *blue* (log_2_-fold enrichment score > 0), while loss-of-function mutations are shown in *red* (log_2_-fold enrichment score ≤ 0). The amino acids of the canonical human PAI-1 sequence are shown in *dark grey*. Amino acids that were either not present or not present at sufficient levels to quantify (BaseMean ≥ 50 as determined by DESeq2 (Love, et al. 2014)) in the phage library are shown in white. (B) Heatmap showing the common amino acid substitutions across PAI-1’s specificity spaces for uPA, FXIIa, and TMPRSS2 with positive (*blue)* and negative (*red*) functional enrichment scores. (C) PCA plot comparing the specificity spaces of PAI-1 for uPA (*pink*), FXIIa (*yellow*), TMPRSS2 (*blue*), and the starting I91L PAI-1 DMS library (*black*). Groups are annotated with the minimum enclosing ellipse. (D and E) Venn diagrams depicting the number of significantly (D) enriched and (E) depleted amino acid substitutions shared and unique to each specificity fingerprint.

### Single amino acid substitutions in PAI-1 differentially affect its ability to inhibit target proteases

We next screened our I91L PAI-1 variant library to identify the effects of single amino acid substitutions in PAI-1 on its ability to inhibit FXIIa and TMPRSS2 in addition to uPA (**Fig. 2A**). Selection with each protease generated a unique PAI-1 variant map (**Fig. 2A**).

The statistically significant functional log_2_-fold enrichment scores are most similar between uPA and FXIIa, although there is a distinct set of mutations that maintain inhibitory activity toward all three proteases (**Fig. 2B**). Similarly, principal component analyses (PCA) of the differential variant enrichment data for PAI-1 specificity towards each of the target proteases (**Fig. 2C**) are most similar for uPA and FXIIa, with TMPRSS2 yielding a relatively distinct pattern of enrichment. The large separation of the principal components required for TMPRSS2 inhibition from that of uPA and FXIIa is consistent with the closer phylogenetic relationship between the latter two proteases (Yousef, et al. 2003) and indicative of more distinct amino acid substitutions required to render PAI-1 a specific/efficient TMPRSS2 inhibitor. We further examined the unique set of amino acid substitutions that render PAI-1 capable (**Fig. 2D**) or incapable (**Fig 2E**) of inhibiting each of these three target proteases using the statistical thresholds described above.

Again, the set of amino acid substitutions in PAI-1 that promote/maintain or result in a loss of uPA or FXIIa inhibition are more similar to each other than either is to those that impact TMPRSS2 inhibition. More amino acid substitutions pass the statistical threshold (p_adj_ < 0.1 and BaseMean score ≥ 50) for uPA inhibition (n = 1369) than for either FXIIa (n = 972) or TMPRSS2 (n = 704) (Table 1), suggesting that the more distantly related a protease is to PAI-1’s canonical target uPA, the fewer possible “pathways” to inhibition exist.

**Table 1.** PAI-1 variants* that lead to maintenance/decrease of inhibitory capacity.

| **Target**<br>**Protease** | **Number of variants that maintain/**<br>**inhibitory capacity** | **Number of variants resulting in**<br>**loss of inhibitory capacity** |
| --- | --- | --- |
| **uPA** | 541 | 828 |
| **FXIIa** | 349 | 623 |
| **TMPRSS2** | 513 | 191 |

- 2 \*Variants identified that pass significance thresholds ( $p_{adj} < 0.1$ , Base Mean score $\geq 50$ ).

The results of our DMS screens of PAI-1 inhibition of three distinct serine protease targets suggest that multiple single amino acid substitutions may result in PAI-1 being a TMPRSS2 inhibitor with impaired or reduced inhibition of its canonical target uPA (**Fig. 2B**). In other words, our results suggest that PAI-1 may exhibit *marginal specificity* with respect to TMPRSS2 inhibition as multiple single amino acid substitutions appear to alter its protease specificity (McClune and Laub 2020; Ghose, et al. 2023). In contrast, the log_2_-fold enrichment scores of the PAI-1 variant library with respect to uPA and FXIIa inhibition closely mirror each other (**Fig. 2**). We hypothesize that this pattern is the result of PAI-1 exhibiting robust specificity for the closely related serine proteases uPA and FXIIa in which single amino acid substitutions do not dramatically alter its protease inhibition profile. However, it remains unclear if PAI-1’s inhibition of FXIIa results from cross-reactivity between two closely related proteins or if PAI-1 is being actively selected to be an inhibitor of both uPA and FXIIa (Yousef, et al. 2003; Conant and Wolfe 2008). In the latter scenario, there is an active selection for PAI-1 to inhibit FXIIa, while in the former, PAI-1 cross-reactivity with FXIIa may be the result of an inherent difficulty in SERPINs evolving insular functional niches with respect to closely related serine proteases (McClune, et al. 2019; McClune and Laub 2020).

### Specificity determinants span PAI-1’s primary structure

To interrogate the role of multiple regions of PAI1 in protease specificity, we next performed separate PCAs on each of the twelve 150 base pair (bp) sequencing amplicons (**Fig. 3**, **SI Table 1**) (Huttinger, et al. 2021; Haynes, et al. 2022). Across all twelve amplicons, principal components (PC) 1 and 2 discriminate between the log_2_-fold enrichment scores with respect to inhibition of uPA, FXIIa, and TMPRSS2, demonstrating that SERPIN regions beyond the RCL (residues 331-350 of human PAI-1 located within amplicon 12) (Gettins and Olson 2009) contribute to target protease specificity. Previous reports of chimeric PAI-1 variants with RCLs from SERPINs targeting alternative proteases retaining the ability to inhibit uPA/tPA are consistent with this finding (Lawrence, et al. 1990).

**Figure 3.**
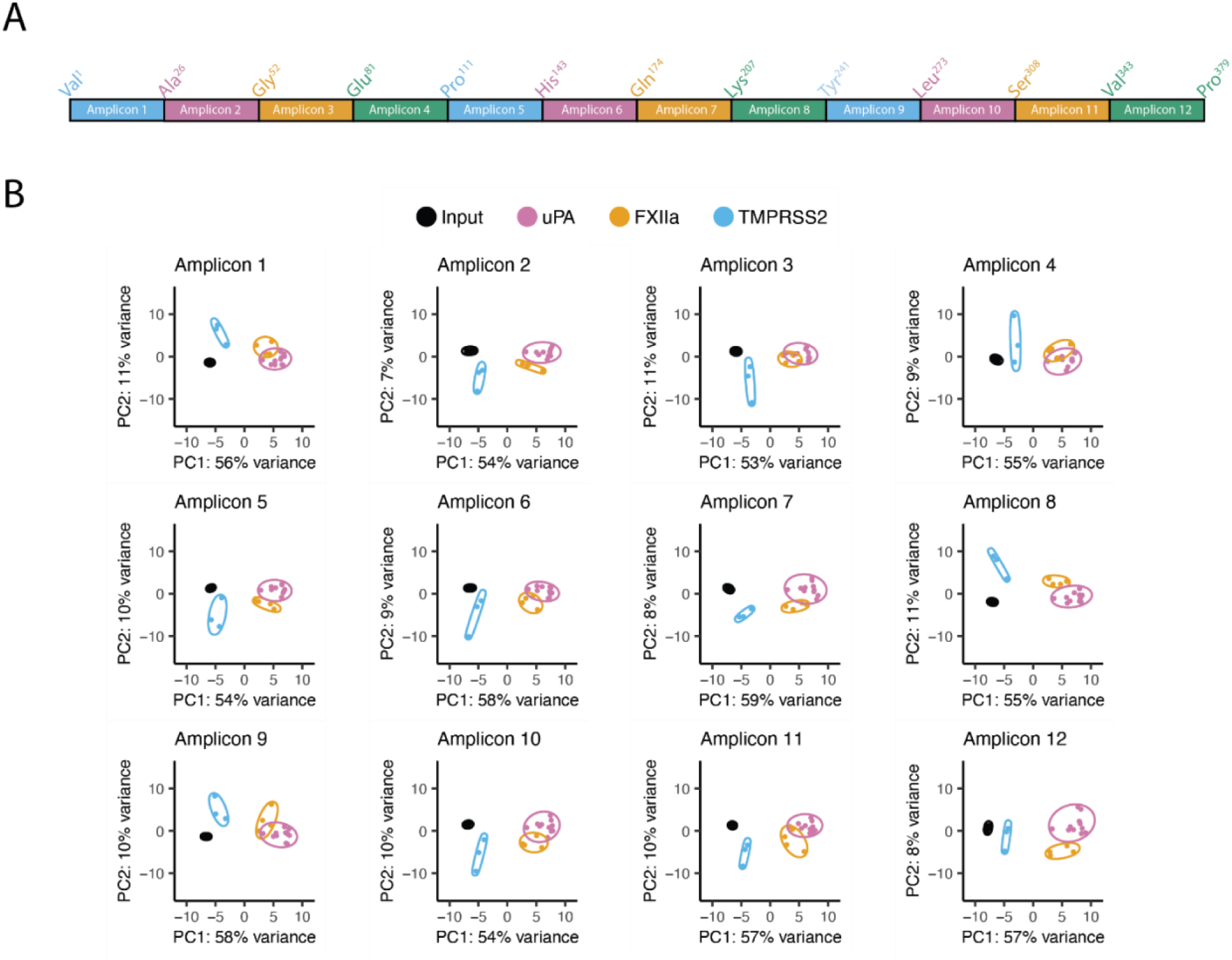
Determinants of PAI-1 specificity are found throughout its sequence space. (A) A cartoon of the PAI-1 primary amino acid sequence showing which amino acids were included in each amplicon. Amplicon 12 contains PAI-1’s RCL. (B) PCA plots comparing the specificity spaces of PAI-1 for uPA (*pink*), FXIIa (*yellow*), TMPRSS2 (*blue*), and the I91L PAI-1 variant library (*black*) across PAI-1’s twelve amplicons. Groups are annotated with the minimum enclosing ellipse.

In contrast, PAI-1 variants with amino acid substitutions in the RCL have been reported to alter PAI-1’s specificity to be either a neutrophil elastase or cathepsin G inhibitor (Stefansson, et al. 2004), similar to the naturally occurring Pittsburgh variant of the SERPIN α_1_-anti-trypsin (A1AT, *SERPINA1*), in which a single amino acid substitution at the P1 position in the RCL converts it from an elastase to a thrombin inhibitor (Lewis, et al. 1978; Owen, et al. 1983), or engineering efforts on other SERPIN backbones to generate SERPINs with novel inhibitory profiles (Scott, et al. 2014; Polderdijk, et al. 2017; Bhakta, et al. 2021; Sanrattana, et al. 2021; Singh, et al. 2022). Therefore, although single amino acid substitutions in a SERPIN’s RCL may function in a manner consistent with marginal specificity for its target protease, our data suggest that multiple amino acid substitutions throughout the SERPIN’s primary sequence may be required to achieve robust specificity for a SERPIN as an efficient and specific inhibitor of a given protease. Furthermore, these variants may contribute to the coevolution of specific SERPIN::serine protease inhibitory reactions through as of yet not fully understood epistatic mechanisms (Spence, et al. 2021; Ding, et al. 2022; Park, et al. 2022) that may improve a SERPIN’s tolerance for efficiently inhibiting one serine protease over another.

### Evolutionary conservation of PAI-1 inhibitory functions

Next, to better understand the evolution of PAI-1’s inhibitory niche, we compared the results of our DMS screens to PAI-1 sequences of extant mammalian species (n = 94). We previously demonstrated that key residues in PAI-1 for inhibition of uPA are conserved in nature by comparing the results of our DMS screen to natural sequence variation across mammals (Huttinger, et al. 2021). We extended these analyses to the present screen to assess whether sites mediating PAI-1 inhibition of FXIIa and TMPRSS2 exhibit similar selective pressures. An evolutionary conservation score for a given site in the PAI-1 amino acid sequence was determined (Ashkenazy, et al. 2016; Huttinger, et al. 2021) and correlated with the tendency of a site to accept mutations that allow PAI-1 to inhibit a target protease (denoted henceforth as the *normalized functional score*). If there is purifying selective pressure for PAI-1 to maintain the inhibition of a given target protease, then we expect that sites in PAI-1 at which amino acid substitutions do not negatively impact the ability to inhibit the target protease will be less conserved across extant species. Likewise, sites at which amino acid substitutions render PAI-1 unable to inhibit the target protease will be more conserved.

The results of these analyses corroborate our earlier findings that PAI-1 inhibition of uPA is under purifying selection (Huttinger, et al. 2021) with a modest, statistically significant correlation between the normalized functional score and evolutionary conservation score across PAI-1 specificity space (slope = 0.58, R^2^ = 0.1427, p = 5.6x10^-14^, **Fig. 4A**). With respect to FXIIa inhibition by PAI-1, the functional and evolutionary conservation scores associate with a smaller slope and weaker, yet statistically significant, correlation (slope = 0.33, R^2^ = 0.066, p = 7.5x10^-7^, **Fig. 4B**). However, no statistically significant association was observed between functional and evolutionary conservation score with respect to PAI-1 inhibition of TMPRSS2 (slope = -0.01, R^2^ = 8.7x10^-5^, p = 0.87, **Fig. 4C**). First, these results suggest that, as expected, PAI-1 inhibition of uPA is under purifying selection. Second, either through FXIIa’s close evolutionary relationship with uPA or via direct effects of selection, sites impacting FXIIa inhibition are also under purifying selection. Meanwhile sites impacting inhibition of TMPRSS2 are not, on average, under differential selection compared to other sites in the molecule. These results suggest that PAI-1 inhibition of TMPRSS2 is potentially an example of cross-functional SERPIN reactivity resulting in inhibition of a non-canonical target and that inhibition of TMPRSS2 is not a biologically significant function of PAI-1 that generates detectable signatures of selection across species.

**Figure 4.**
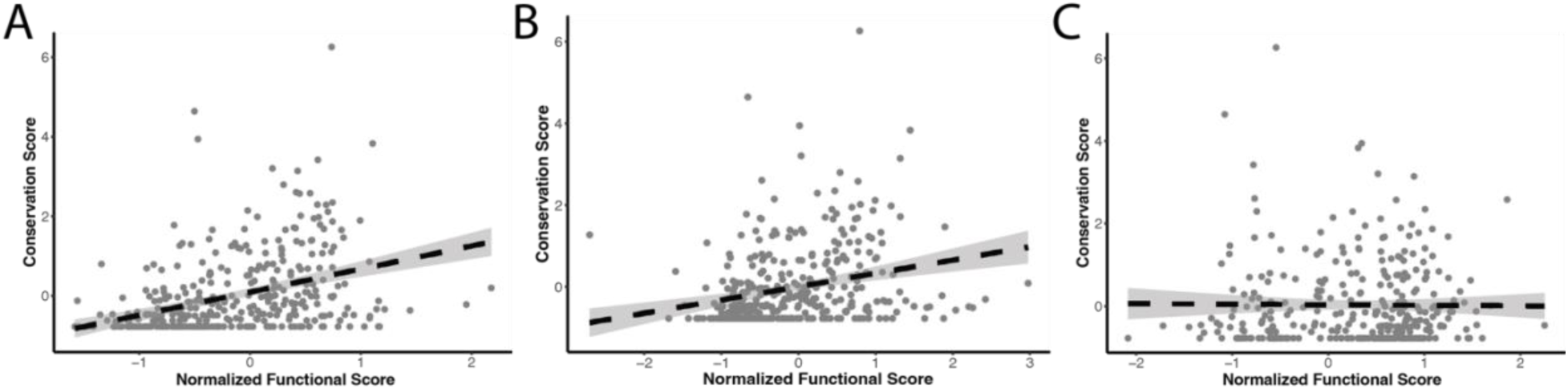
PAI-1 is under purifying selection to inhibit uPA and FXIIa but not TMPRSS2. Conservation scores (Ashkenazy, et al. 2016) for each amino acid position in PAI-1 (*grey dots*) are shown as a function of the normalized functional scores determined in our DMS screens of PAI-1 inhibition of (A) uPA (R^2^ = 0.14, p = 4.6x10^-14^), (B) FXIIa (R^2^ = 0.06, p = 7.5x10^-7^), and (C) TMPRSS2 (R^2^ = -0.003, p = 0.87). Dashed lines indicated the best fit linear regression with the 95% confidence interval shown in *grey* shading.

Given that a greater proportion of PAI-1 sequence variation across extant species is explained by a drive to maintain PAI-1’s ability to inhibit uPA than FXIIa, we speculate that FXIIa inhibition by PAI-1 may be the result of cross-reactivity between uPA and FXIIa, given their close phylogenetic relationship, while TMPRSS2 represents a distinct outgroup to the uPA/FXIIa clade (Yousef, et al. 2003). The *F12* gene appeared in stem tetrapods after their divergence from fish, and therefore interactions between PAI-1 and uPA predate the existence of FXIIa (Ponczek, et al. 2008; Ponczek, et al. 2020). Insulation of PAI-1 from crosstalk to FXIIa may have been prevented by the evolutionary inertia of millions of years of preceding coevolution between PAI-1 and plasminogen activators.

### Comparing the effects of single amino acid substitution in PAI-1 on the inhibition of uPA and FXIIa

Although PAI-1 inhibits both uPA and FXIIa, PAI-1 inhibition of uPA is anti-fibrinolytic (stabilizing the thrombus) while inhibition of FXIIa is anti-thrombotic. One potential explanation for these opposite effects, is that due to the recent divergence of uPA and FXIIa, PAI-1’s ability to inhibit uPA has been “carried over” into its ability to inhibit FXIIa, as evidenced by the similarity in PAI-1’s specificity fingerprints for both uPA and FXIIa (**Figs. 2** and **5**). To further address the similar effects of amino acid substitution in PAI-1 on its ability to inhibit uPA and FXIIa, we next tested the effects of the most enriched amino acid substitutions in PAI-1 with respect to the inhibition of both of these proteases. We identified the most enriched single amino acid substitutions in our DMS screens for PAI-1 inhibition of both uPA and FXIIa on the I91L background as being F98Y, T251I, and S331C (**Fig. 5A** and **5B**). The WT human amino acids at these positions are highly conserved in extant mammalian species, possibly due to the requirement of the I91L substitution or a general extension of PAI-1’s functional half-life to open up the functional accessibility of these substitutions.

**Figure 5.**
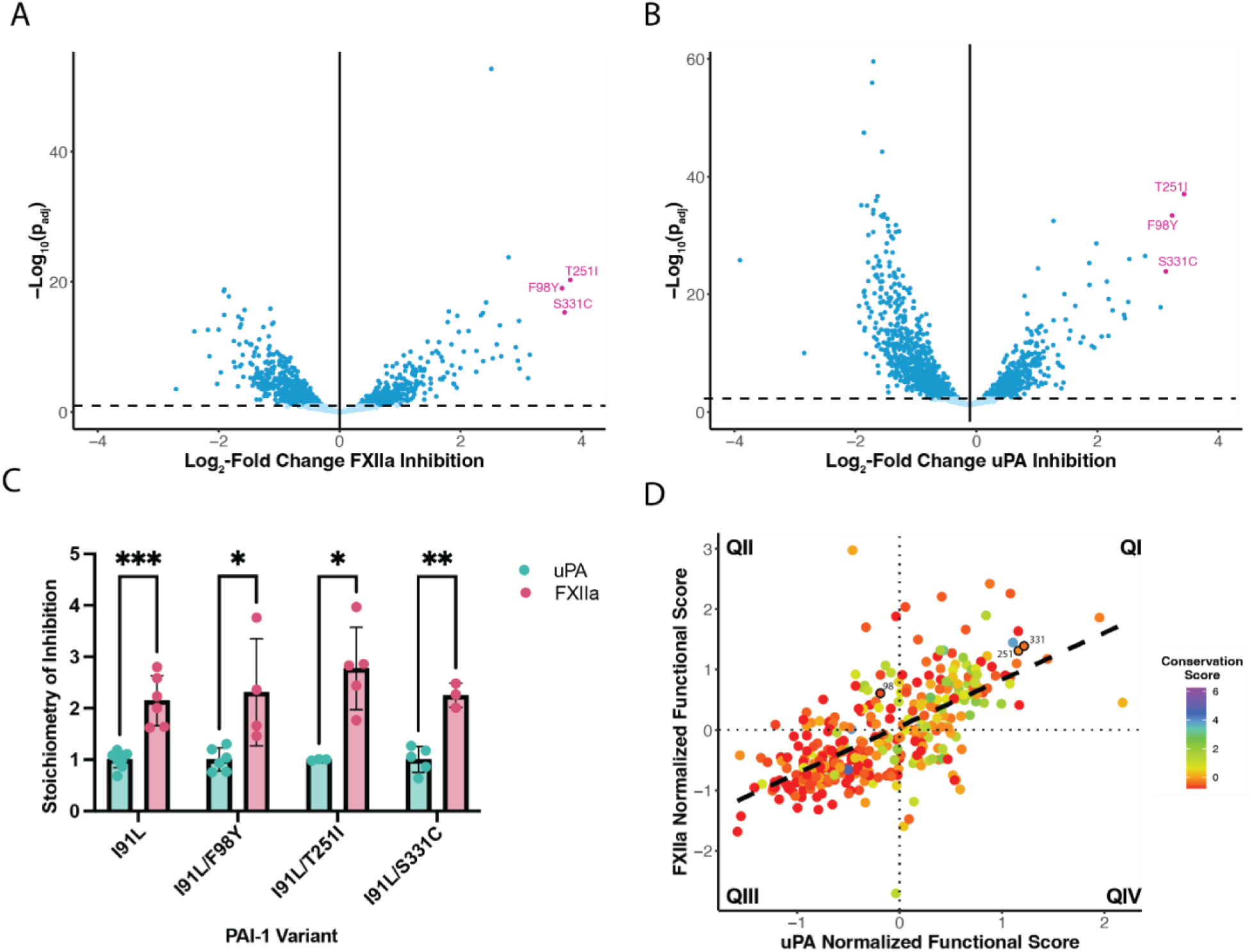
PAI-1 variants that are most enriched for FXIIa inhibition do not exhibit improved SIs. Volcano plots showing the I91L PAI-1 variants activity toward (A) FXIIa and (B) uPA. Dark blue circles indicate statistically significant enrichment scores (p < 0.005). The three most enriched PAI-1 variants (I91L/F98Y, I91L/T251I, and I91L/S331C) are shown in pink. The Log_2_-fold change is plotted on the x-axis and the statistical significance (-Log_10_(p_adj_)) is plotted on the y-axis. (C) SIs were determined for each of the variants for both uPA (green) and FXIIa (pink) inhibition. For each variant, the SI for uPA was set to one to account for inactive PAI-1 present in the protein preparation. t-tests were used to determine statistical significance (*, p <0.05; **, p<0.01; ***, p < 0.001). (D) FXIIa normalized functional mutation scores are compared to the uPA normalized functional mutation scores for each variant. The conservation score is indicated by the color of the points. Residues 98, 251, and 331 are labelled and outlined in black.

These variants were expressed as recombinant proteins, and their SIs with respect to uPA and FXIIa were determined (**Fig. 5C**). For a SERPIN to function as a specific and efficient inhibitor of a given serine protease, it must have a sufficiently rapid second-order rate constant and a SI close to “1” such that virtually all the reactions proceed along the inhibitory pathway with little to no substrate pathway observed (Olson and Gettins 2011). The SI for FXIIa inhibition was consistently 2-3 fold higher than that of uPA inhibition—indicating that when interacting with FXIIa, in contrast to uPA, PAI-1 is more likely to utilize the substrate versus the inhibitory pathway (**Fig. 1**). Of note, these variants were not the most enriched variants in our original DMS screen on the WT PAI-1 background (Huttinger, et al. 2021), with the F98Y and S331C variants actually classified as non-functional against WT PAI-1 (**SI Fig. 2C**). This finding suggests that these mutations are epistatic with the I91L substitution with respect to PAI-1 function (Berkenpas, et al. 1995) or that these variants decrease the functional stability of PAI-1 such that their half-lives are too short to be detected on the WT background in our assay, despite being otherwise inhibitory variants. One potential implication of epistasis between the I91L and T98Y and/or S331C substitutions, with respect to PAI-1 inhibitory function, is that although the T98Y and S331C substitutions are functional inhibitors on the I91L background, this functionality of these substitutions is not available to PAI-1 in the absence of the I91L substitution.

As the three most highly enriched PAI-1 variants identified in our DMS screens for both uPA and FXIIa inhibition were the same (**Figs. 5A** and **5B**), we next compared the evolutionary conservation scores for PAI-1 with respect to both uPA and FXIIa inhibition (**Fig. 5D**). Not only are the normalized functional scores for uPA and FXIIa highly correlated with each other (slope = 0.78, R^2^ = 0.43, p = 2.2x10^-16^) as expected (**Fig. 2B**); but amino acid sites that are conserved across extant mammalian species are less likely to accept mutations in both DMS screens (quadrant I, QI), while sites that are more evolutionary labile are more likely to accept mutations (quadrant III, QIII). The mean evolutionary conservation scores (± standard deviation) between Q1 (0.45 ± 1.21) and QIII (-0.36 ± 0.70) are highly statistically significantly different (p_adj_ < 0.0001, Tukey’s HSD test), suggesting that amino acid substitutions in PAI-1 that would impact its protease specificity for either uPA or FXIIa are likely to affect inhibition of both proteases.

It has been speculated that the high SI for PAI-1 inhibition of FXIIa effectively results in FXIIa inactivation of PAI-1, further reducing its potential as an antifibrinolytic (Tanaka, et al. 2009). As the inhibition of both uPA and FXIIa occupy similar PAI-1 sequence spaces, the amino acid substitutions that were identified as the best FXIIa inhibitors in our DMS screen were still more efficient uPA inhibitors (**Fig. 5**)—consistent with previous studies (Berrettini, et al. 1989; Sherman, et al. 1992). Overall, our data indicate that PAI-1 maintains its function as an inhibitor of these two proteases (although there is a “preference” to inhibit uPA over FXIIa), with few single amino acid substitutions (n = 16) conferring a preference for inhibition of FXIIa over uPA (**Figs. 2B** and **5A-B**). This is in contrast to PAI-1 inhibition of TMPRSS2, in which multiple single amino acid substitutions (n = 184) appear to improve specificity toward TMPRSS2 over FXIIa/uPA(**Fig. 2B**). We, therefore, hypothesized that PAI-1 has maintained inhibitory activity toward FXIIa in addition to uPA as a function of the close evolutionary relationship between these serine proteases(Yousef, et al. 2003; McClune, et al. 2019). In essence, our results suggest closely related serine proteases form a crowded sequence space with regard to SERPIN inhibition. An expectation arising from these findings is that, beyond PAI-1, a given SERPIN will inhibit groups of closely related serine proteases and patterns of SERPIN and protease diversification may mirror one another.

### Coevolution of serpins and their cognate serine proteases

To test the role of phylogenetic relatedness and codiversification in driving interactions between SERPINs and their cognate serine proteases, we next performed a co-phylogenetic analysis of human secreted SERPINs and their cognate serine proteases. Cophylogenetic analyses are typically used to study coevolution in host-parasite interactions, where the null hypothesis is that the shapes of the two phylogenies of each putatively coevolving group are not mutually informative of one another (Legendre, et al. 2002). We reasoned that a similar statistic of mutual information in phylogenetic topologies would quantify SERPIN:serine protease codiversification. We therefore curated a list of human secreted SERPINs (n = 13) with well-described cognate serine proteases (n = 16) (Olson and Gettins 2011; Heiker, et al. 2013; Godinez, et al. 2022; Janciauskiene, et al. 2024).

Our cophylogenetic analysis (**Fig. 6**) supports the hypothesis that the secreted SERPINs and serine proteases have co-diversified (p = 0.039). The bulk of this mutual signal appears to reside deep in the evolution of these groups, reflected in the SERPINA clade largely inhibiting the KLK/elastase clade, whereas the other SERPINs, such as the clade formed by group I/E/G SERPINs, largely inhibit the related complement system and coagulation proteases. Notably for the current study, the closely related *SERPINE1*, *SERPINE2*, and *SERPINI1* all inhibit uPA (*PLAU*) and tPA (*PLAT*) and most likely coevolved from an ancestral SERPIN, with the emergence of the two plasminogen activators from a common serine protease ancestor that also gave rise to FXIIa (*F12*) (Jendroszek, et al. 2019). Similarly, the closely related serine proteases thrombin (encoded by the *F2* gene) and FXa (encoded by the *F10* gene) are both inhibited by a common SERPIN, antithrombin (encoded by the *SERPINC1* gene), with which they also coevolved (**Fig. 7**) (Yousef, et al. 2003). Therefore, we anticipate that a DMS screen of antithrombin’s inhibition of thrombin and FXa would also reveal overlapping effects of single amino acid substitutions.

**Figure 6.**
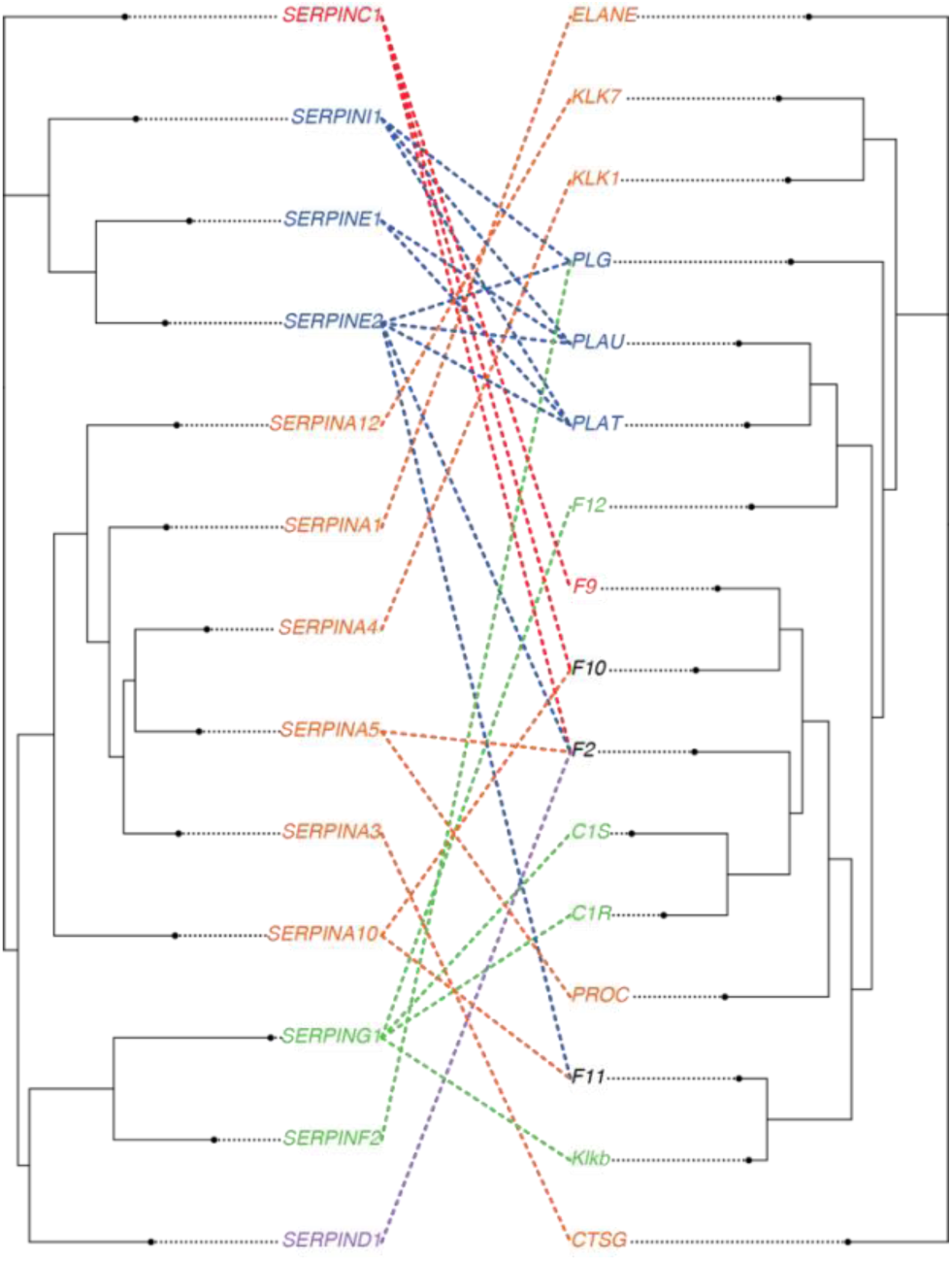
Cophylogenetic analysis reveals that human secreted SERPINs coevolved with secreted serine proteases. The phylogenic tree of human secreted SERPINs (*left*) is mutually informative with that of human secreted serine proteases (*right*) as determined with the parafit R package (p = 0.039) (Legendre, et al. 2002). SERPINs and proteases are denoted by their human gene names. For SERPINs, colors denote closely related proteins (SERPINC clade, red; SERPIN E and I clades, blue; SERPIN A clade, orange; SERPIN G and F clades, green; SERPIN D clade, purple), while for serine proteases, colors denote the group of SERPINs most commonly inhibiting a given protease. Proteases equally likely to be inhibited by SERPINs from more than one group are shown in black.

## Conclusion

Inhibitory SERPINs have evolved to fulfill unique functional niches in which they inhibit one or a few proteases with residual inhibitory activity towards additional proteases (Gettins 2002; Olson and Gettins 2011). Here, we have harnessed the power of DMS and combined it with comparative evolutionary approaches to assess PAI-1’s inhibitory function towards its canonical target, uPA, as well as non-canonical targets that are both closely (FXIIa) and distantly (TMPRSS2) related to uPA. Our data demonstrate that the determinants of PAI-1 specificity are scattered throughout its amino acid sequence space, with no one region of the primary sequence completely dictating the specificity of a target protease, suggesting that residues within the PAI-1 structure mediate long-range effects that help dictate its specificity as a protease inhibitor. Furthermore, we find that residues modulating PAI-1 inhibition of uPA and FXIIa are under purifying selection, with no such selective signature for inhibition of TMPRSS2. PAI-1 instead exhibits marginal specificity towards TMPRSS2, but robust specificity that prevents it from becoming a FXIIa specialist. As uPA and FXIIa are closely related serine proteases, the ability of PAI-1 to be a specific inhibitor of uPA plasminogen activators, but not FXIIa, is restricted by a locally crowded sequence space. Coevolution of SERPINs with their cognate target serine protease may explain many overlapping and non-specific SERPIN::serine protease interactions, and could help guide future efforts to engineer SERPIN specificity. In conclusion, we have demonstrated the power of DMS to interpret the evolution and functional specificity of members of the SERPIN protein superfamily and anticipate that similar analyses have the potential to improve understanding of the mechanisms driving the diversification and specialization of other large paralogous protein families.

## Materials and Methods

### Phage displayed PAI-1 variant library preparation

A phage display PAI-1 variant library was constructed on the I91L PAI-1 background using error prone PCR as previously described (7, 8) with the GeneMorph II Random Mutagenesis Kit (Agilent Technologies, Santa Clara, CA) using primers that preserved the AscI and NotI restriction sites, and cloned into a modified pAY-FE plasmid (Genebank #MW464120) in which an amber stop codon (*TAG*) immediately preceding the SERPINE1-gIII fusion construct was mutated to glutamine (CAG, Gln) to increase expression of the PAI-1-pIII coat fusion protein (pLMH1, **SI Table 1**) (Huttinger, et al. 2021; Haynes, et al. 2022). Following cloning of the I91L PAI-1 variant library into the pLMH1 plasmid, the library was transformed into electrocompetent XL-1 Blue MRF’ *E. coli* (Agilent Technologies). Library depth was determined by counting the number of ampicillin-resistant colonies. Mutation frequency was estimated by Sanger sequencing of the PAI-1 (*SERPINE1*) inserts from randomly selected colonies (n=24).

The I19L PAI-1 phage displayed library was produced as described previously (Huttinger, et al. 2021; Haynes, et al. 2022). Briefly, *E. coli* harboring the I91L PAI-1 library were grown in LB media supplemented with 2% glucose and ampicillin (0.1 mg/mL) to mid-log phase at 37°C, infected with M13KO7 helper phage (Cytiva) for 1 h, and transferred to 2xYT media supplemented with ampicillin (0.1 mg/mL), kanamycin (0.03 mg/mL) and IPTG (0.4 mM) to induce expression of the PAI-1 phage displayed library and grown for 2h at 37°C. *E. coli* were removed by centrifugation (4200xg and 4500xg at 4°C for 10 min) and phage were precipitated from the supernatant with PEG-8000 (2.5% w/v) and NaCl (0.5 M) overnight at 4°C. Phage were pelleted by centrifugation (20,000xg for 20 min at 4°C) and resuspended in 50 mM Tris containing 150 mM NaCl at pH 7.4 (TBS).

### PAI-1 variant selection

uPA selection assays were performed as previously described (Huttinger, et al. 2021; Haynes, et al. 2022). Human coagulation FXIIa (Innovative Research, Novi, MI) was biotinylated (FXIIa-biotin) and the degree of labeling assessed using the EZ-Link Sulfo-NHS-LC-Biotin labelling kit (ThermoFisher Scientific) with 1-2 molecules of biotin per protein molecule. Soluble recombinant TMPRSS2 (aa 106-492) with an N-terminal 6xHis tag expressed in yeast was purchased from Creative BioMart (Shirley, NY) and shown to be inhibited by WT PAI-1 (**SI Fig. 1**). FXIIa-biotin (100 nM) or TMPRSS2 (1 uM (Shrimp, et al. 2020)) were incubated with a 1:10 dilution of the input I91L PAI-1 phage display library for 30 min at 37°C in TBS (50 mM Tris base, 150 mM NaCl, pH 7.4) containing 5% BSA (TBS-BSA). All reactions were quenched with the addition of cOmplete EDTA-free protease inhibitor cocktail (MilliporeSigma) for 10 min at 37°C. The biotinylated and 6xHis-tagged screens were immunoprecipitated overnight with streptavidin or Ni-NTA magnetic beads (New England Biolabs, Ipswich, MA), respectively. phPAI-1::enzyme complexes were further washed, eluted, and quantified as described previously (Huttinger, et al. 2021; Haynes, et al. 2022). Replicas represent the selection of independent cultures of the input phage display PAI1 variant library.

### HTS

Phage displayed PAI-1 cDNA were sequenced in twelve 150 bp amplicons using primers listed in **SI Table 1**. Amplicons were prepared for HTS as previously described (Huttinger, et al. 2021; Haynes, et al. 2022) and sequenced with 8x10^5^-2x10^6^ 150 bp paired-end reads per amplicon. HTS data were analyzed using the DESeq2 software package (Love, et al. 2014; Zhu, et al. 2019). Significance thresholds for DESeq2 were set to p_adj_ < 0.1 and BaseMean score ≥ 50. MA plots showing the distribution of amino acid substitutions in PAI-1 with respect to these thresholds are shown in **SI Figs. 2-4**. Notably, when amber stop codons are read through as Gln, those variants are enriched in the selected libraries relative to other stop codons introduced by error-prone PCR.

### Evolutionary variability of PAI-1

As previously described, evolutionary conservation scores at each amino acid position in PAI-1 were calculated using ConSurf (https://consurf.tau.ac.il/), where higher ConSurf scores indicate more evolutionarily variable positions (Ashkenazy, et al. 2016; Huttinger, et al. 2021). To relate the results of our DMS screen to evolutionary conservation scores, we define the *normalized functional score* as the sum of the mean log_2_-fold determined at each position. Conservation scores were then analyzed as a function of the normalized functional score at each position using the R software package (version 4.3.3).

### Expression and characterization of recombinant PAI-1 variants

PAI-1 variant cDNA (I91L, I91L/F98Y, I91L/T251I, and I91L/S331C PAI-1) in the pET-24(+) expression plasmid with a C-terminal Gly-Ser-Gly hinge and 6XHis-tag was purchased from Twist Bioscience (South San Francisco, CA) and transformed into NiCo21(DE3) chemically competent *E. coli* (New England Biolabs, Ipswitch, MA). PAI-1 variants were expressed as previously described (Haynes, et al. 2022), and concentration was determined by absorption at 280 nm (ε_1%, 280 nm_ = 7.0). SI for each variant was determined by incubating uPA (2.5 nM) or factor XIIa (100 nM) with PAI-1 variant concentrations ranging from 0-4 nM and 0-10 nM respectively. The SIs for each variant were normalized to an SI of uPA inhibition defined as 1.

### PAI-1 inhibition of TMPRSS2

WT PAI-1 was expressed and purified as described above. TMPRSS2 (1 μM total protein, yet only fractionally active (Shrimp, et al. 2020)) was incubated with WT PAI-1 (1 nM effective active concentration) or vehicle control for 30 min at room temperature (∼25°C). TMPRSS2 activity was determined by monitoring its ability to cleave peptidyl substrate boc-QAR-AMC (λ_ex_ = 370 nm, λ_em_ = 440 nm; VWR International) for 10 min (Shrimp, et al. 2020). Following this monitoring, samples were run on a 4-20% Tris-glycine gel (Invitrogen) under non-reducing conditions, transferred to nitrocellulose, and blotted using a rabbit-anti-PAI-1 primary antibody (Abcam).

### Cophylogenetic analyses

A list of human secreted SERPINs (n = 13) and serine proteases (n = 16) was curated from the literature (Olson and Gettins 2011; Heiker, et al. 2013; Godinez, et al. 2022; Janciauskiene, et al. 2024). We defined secreted SERPINs and serine proteases as those containing a signal peptide. We used the COBALT tool to produce a domain-centric alignment of both the SERPINs and serine proteases in the lists, and then pruned the alignment manually to include only sites with fewer than 90% gaps. We used TreeBeST (Vilella, et al. 2009) to produce phylogenetic trees for both SERPINs and serine proteases, using 1000 bootstrap replicates, and recovered the maximum likelihood tree for analyses. A cophylogenetic analysis was then performed using the Parafit package in R with the null hypothesis that the evolution of serine proteases and SERPINS were independent of each other (Legendre, et al. 2002). The p-value was derived by 1000 random permutations of the tip states of the two trees.

## Data Availability

All data is contained within the manuscript and supplementary files. Raw sequencing data and bioinformatics results will be made available upon request.

## References

Al-Samkari H, Karp Leaf RS, Dzik WH, Carlson JCT, Fogerty AE, Waheed A, Goodarzi K, Bendapudi PK, Bornikova L, Gupta S, et al. 2020. COVID-19 and coagulation: bleeding and thrombotic manifestations of SARS-CoV-2 infection. Blood 136:489–500.

Ashkenazy H, Abadi S, Martz E, Chay O, Mayrose I, Pupko T, Ben-Tal N. 2016. ConSurf 2016: an improved methodology to estimate and visualize evolutionary conservation in macromolecules. Nucleic Acids Res 44:W344–350.

Berkenpas MB, Lawrence DA, Ginsburg D. 1995. Molecular evolution of plasminogen activator inhibitor-1 functional stability. EMBO J 14:2969–2977.

Berrettini M, Schleef RR, España F, Loskutoff DJ, Griffin JH. 1989. Interaction of type 1 plasminogen activator inhibitor with the enzymes of the contact activation system. Journal of Biological Chemistry 264:11738–11743.

Bhakta V, Hamada M, Nouanesengsy A, Lapierre J, Perruzza DL, Sheffield WP. 2021. Identification of an alpha-1 antitrypsin variant with enhanced specificity for factor XIa by phage display, bacterial expression, and combinatorial mutagenesis. Sci Rep 11:5565.

Bouwman JJ, Diepersloot RJ, Visseren FL. 2009. Intracellular infections enhance interleukin-6 and plasminogen activator inhibitor 1 production by cocultivated human adipocytes and THP-1 monocytes. Clin Vaccine Immunol 16:1222–1227.

Conant GC, Wolfe KH. 2008. Turning a hobby into a job: how duplicated genes find new functions. Nat Rev Genet 9:938–950.

Ding D, Green AG, Wang B, Lite TV, Weinstein EN, Marks DS, Laub MT. 2022. Co-evolution of interacting proteins through non-contacting and non-specific mutations. Nat Ecol Evol 6:590–603.

Dittmann M, Hoffmann HH, Scull MA, Gilmore RH, Bell KL, Ciancanelli M, Wilson SJ, Crotta S, Yu Y, Flatley B, et al. 2015. A serpin shapes the extracellular environment to prevent influenza A virus maturation. Cell 160:631–643.

Gettins PG. 2002. Serpin structure, mechanism, and function. Chem Rev 102:4751–4804.

Gettins PG, Olson ST. 2009. Exosite determinants of serpin specificity. J Biol Chem 284:20441–20445.

Ghose DA, Przydzial KE, Mahoney EM, Keating AE, Laub MT. 2023. Marginal specificity in protein interactions constrains evolution of a paralogous family. Proc Natl Acad Sci U S A 120:e2221163120.

Godinez A, Rajput R, Chitranshi N, Gupta V, Basavarajappa D, Sharma S, You Y, Pushpitha K, Dhiman K, Mirzaei M, et al. 2022. Neuroserpin, a crucial regulator for axogenesis, synaptic modelling and cell-cell interactions in the pathophysiology of neurological disease. Cell Mol Life Sci 79:172.

Hahn MW. 2009. Distinguishing among evolutionary models for the maintenance of gene duplicates. J Hered 100:605–617.

Haynes LM, Huttinger ZM, Yee A, Kretz CA, Siemieniak DR, Lawrence DA, Ginsburg D. 2022. Deep mutational scanning and massively parallel kinetics of plasminogen activator inhibitor-1 functional stability to probe its latency transition. J Biol Chem 298:102608.

Heiker JT, Klöting N, Kovacs P, Kuettner EB, Sträter N, Schultz S, Kern M, Stumvoll M, Blüher M, Beck-Sickinger AG. 2013. Vaspin inhibits kallikrein 7 by serpin mechanism. Cell Mol Life Sci 70:2569–2583.

Huntington JA. 2011. Serpin structure, function and dysfunction. J. Thromb. Haemost. 9 Suppl 1:26–34.

Huttinger ZM, Haynes LM, Yee A, Kretz CA, Holding ML, Siemieniak DR, Lawrence DA, Ginsburg D. 2021. Deep mutational scanning of the plasminogen activator inhibitor-1 functional landscape. Sci Rep 11:18827.

Janciauskiene S, Lechowicz U, Pelc M, Olejnicka B, Chorostowska-Wynimko J. 2024. Diagnostic and therapeutic value of human serpin family proteins. Biomed Pharmacother 175:116618.

Jendroszek A, Madsen JB, Chana-Munoz A, Dupont DM, Christensen A, Panitz F, Fuchtbauer EM, Lovell SC, Jensen JK. 2019. Biochemical and structural analyses suggest that plasminogen activators coevolved with their cognate protein substrates and inhibitors. J Biol Chem 294:3794–3805.

Keijer J, Linders M, Wegman JJ, Ehrlich HJ, Mertens K, Pannekoek H. 1991. On the Target Specificity of Plasminogen Activator Inhibitor 1: The Role of Heparin, Vitronectin, and the Reactive Site. Blood 78:1254–1261.

Keller TT, Sluijs KFvd, Kruif MDd, Gerdes VEA, Meijers JCM, Florquin S, Poll Tvd, Gorp ECMv, Brandjes DPM, Büller HR, et al. 2006. Effects on Coagulation and Fibrinolysis Induced by Influenza in Mice With a Reduced Capacity to Generate Activated Protein C and a Deficiency in Plasminogen Activator Inhibitor Type 1. Circulation Research 99:1261–1269.

Law RH, Zhang Q, McGowan S, Buckle AM, Silverman GA, Wong W, Rosado CJ, Langendorf CG, Pike RN, Bird PI, et al. 2006. An overview of the serpin superfamily. Genome Biol 7:216.

Lawrence DA, Ginsburg D, Day DE, Berkenpas MB, Verhamme IM, Kvassman JO, Shore JD. 1995. Serpin-protease complexes are trapped as stable acyl-enzyme intermediates. J. Biol. Chem. 270:25309–25312.

Lawrence DA, Olson ST, Muhammad S, Day DE, Kvassman JO, Ginsburg D, Shore JD. 2000. Partitioning of serpin-proteinase reactions between stable inhibition and substrate cleavage is regulated by the rate of serpin reactive center loop insertion into beta-sheet A. J. Biol. Chem. 275:5839–5844.

Lawrence DA, Strandberg L, Ericson J, Ny T. 1990. Structure-function studies of the SERPIN plasminogen activator inhibitor type 1. Analysis of chimeric strained loop mutants. J. Biol. Chem. 265:20293–20301.

Legendre P, Desdevises Y, Bazin E. 2002. A statistical test for host-parasite coevolution. Syst Biol 51:217–234.

Lewis JH, Iammarino RM, Spero JA, Hasiba U. 1978. Antithrombin Pittsburgh: an alpha1-antitrypsin variant causing hemorrhagic disease. Blood 51:129–137.

Love MI, Huber W, Anders S. 2014. Moderated estimation of fold change and dispersion for RNA-seq data with DESeq2. Genome Biol 15:550.

Marijanovic EM, Fodor J, Riley BT, Porebski BT, Costa MGS, Kass I, Hoke DE, McGowan S, Buckle AM. 2019. Reactive centre loop dynamics and serpin specificity. Sci Rep 9:3870.

McClune CJ, Alvarez-Buylla A, Voigt CA, Laub MT. 2019. Engineering orthogonal signalling pathways reveals the sparse occupancy of sequence space. Nature 574:702–706.

McClune CJ, Laub MT. 2020. Constraints on the expansion of paralogous protein families. Curr Biol 30:R460–r464.

Nocedal I, Laub MT. 2022. Ancestral reconstruction of duplicated signaling proteins reveals the evolution of signaling specificity. eLife 11.

Olson ST, Gettins PG. 2011. Regulation of proteases by protein inhibitors of the serpin superfamily. Prog Mol Biol Transl Sci 99:185–240.

Owen MC, Brennan SO, Lewis JH, Carrell RW. 1983. Mutation of antitrypsin to antithrombin. alpha 1-antitrypsin Pittsburgh (358 Met leads to Arg), a fatal bleeding disorder. N. Engl. J. Med. 309:694–698.

Park Y, Metzger BPH, Thornton JW. 2022. Epistatic drift causes gradual decay of predictability in protein evolution. Science 376:823–830.

Polderdijk SG, Adams TE, Ivanciu L, Camire RM, Baglin TP, Huntington JA. 2017. Design and characterization of an APC-specific serpin for the treatment of hemophilia. Blood 129:105–113.

Ponczek MB, Gailani D, Doolittle RF. 2008. Evolution of the contact phase of vertebrate blood coagulation. J Thromb Haemost 6:1876–1883.

Ponczek MB, Shamanaev A, LaPlace A, Dickeson SK, Srivastava P, Sun MF, Gruber A, Kastrup C, Emsley J, Gailani D. 2020. The evolution of factor XI and the kallikrein-kinin system. Blood Adv 4:6135–6147.

Puy C, Ngo ATP, Pang J, Keshari RS, Hagen MW, Hinds MT, Gailani D, Gruber A, Lupu F, McCarty OJT. 2019. Endothelial PAI-1 (Plasminogen Activator Inhibitor-1) Blocks the Intrinsic Pathway of Coagulation, Inducing the Clearance and Degradation of FXIa (Activated Factor XI). Arterioscler Thromb Vasc Biol 39:1390–1401.

Rezaie AR. 2001. Vitronectin Functions as a Cofactor for Rapid Inhibition of Activated Protein C by Plasminogen Activator Inhibitor-1: Implications for the mechanism of profibrinolytic action of activated protein C. J Biol Chem 276:15567–15570.

Sanrattana W, Maas C, de Maat S. 2019. SERPINs—From Trap to Treatment. Front Med (Lausanne) 6:25.

Sanrattana W, Sefiane T, Smits S, van Kleef ND, Fens MH, Lenting PJ, Maas C, de Maat S. 2021. A reactive center loop-based prediction platform to enhance the design of therapeutic SERPINs. Proc Natl Acad Sci U S A 118.

Scott BM, Matochko WL, Gierczak RF, Bhakta V, Derda R, Sheffield WP. 2014. Phage display of the serpin alpha-1 proteinase inhibitor randomized at consecutive residues in the reactive centre loop and biopanned with or without thrombin. PLoS One 9:e84491.

Sen P, Komissarov AA, Florova G, Idell S, Pendurthi UR, Vijaya Mohan Rao L. 2011. Plasminogen activator inhibitor-1 inhibits factor VIIa bound to tissue factor. J Thromb Haemost 9:531–539.

Shen LW, Mao HJ, Wu YL, Tanaka Y, Zhang W. 2017. TMPRSS2: A potential target for treatment of influenza virus and coronavirus infections. Biochimie 142:1–10.

Sherman PM, Lawrence DA, Yang AY, Vandenberg ET, Paielli D, Olson ST, Shore JD, Ginsburg D. 1992. Saturation mutagenesis of the plasminogen activator inhibitor-1 reactive center. J. Biol. Chem. 267:7588–7595.

Shrimp JH, Kales SC, Sanderson PE, Simeonov A, Shen M, Hall MD. 2020. An Enzymatic TMPRSS2 Assay for Assessment of Clinical Candidates and Discovery of Inhibitors as Potential Treatment of COVID-19. ACS Pharmacol Transl Sci 3:997–1007.

Singh S, O’Reilly S, Gewaid H, Bowie AG, Gautier V, Worrall DM. 2022. Reactive Centre Loop Mutagenesis of SerpinB3 to Target TMPRSS2 and Furin: Inhibition of SARS-CoV-2 Cell Entry and Replication. International Journal of Molecular Sciences 23:12522.

Spence MA, Mortimer MD, Buckle AM, Minh BQ, Jackson CJ. 2021. A Comprehensive Phylogenetic Analysis of the Serpin Superfamily. Mol Biol Evol 38:2915–2929.

Stefansson S, Yepes M, Gorlatova N, Day DE, Moore EG, Zabaleta A, McMahon GA, Lawrence DA. 2004. Mutants of plasminogen activator inhibitor-1 designed to inhibit neutrophil elastase and cathepsin G are more effective in vivo than their endogenous inhibitors. J Biol Chem 279:29981–29987.

Tanaka A, Suzuki Y, Sugihara K, Kanayama N, Urano T. 2009. Inactivation of plasminogen activator inhibitor type 1 by activated factor XII plays a role in the enhancement of fibrinolysis by contact factors in-vitro. Life Sci 85:220–225.

Vilella AJ, Severin J, Ureta-Vidal A, Heng L, Durbin R, Birney E. 2009. EnsemblCompara GeneTrees: Complete, duplication-aware phylogenetic trees in vertebrates. Genome Res 19:327–335.

Yousef GM, Kopolovic AD, Elliott MB, Diamandis EP. 2003. Genomic overview of serine proteases. Biochemical and Biophysical Research Communications 305:28–36.

Zhu A, Ibrahim JG, Love MI. 2019. Heavy-tailed prior distributions for sequence count data: removing the noise and preserving large differences. Bioinformatics 35:2084–2092.

Zuo Y, Warnock M, Harbaugh A, Yalavarthi S, Gockman K, Zuo M, Madison JA, Knight JS, Kanthi Y, Lawrence DA. 2021. Plasma tissue plasminogen activator and plasminogen activator inhibitor-1 in hospitalized COVID-19 patients. Sci Rep 11:1580.

