## Supplementary material for "High-throughput amino acid-level characterization of the interactions of plasminogen activator inhibitor-1 with variably divergent proteases": SI Fig. 1

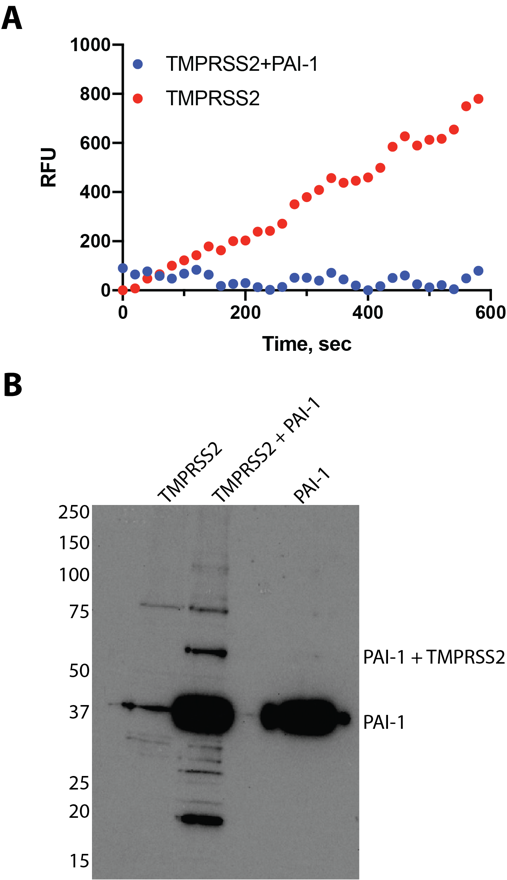

**SI Fig. 1. Direct evidence that PAI-1 inhibits TMPRSS2.** (A) Recombinant soluble TMPRSS2 (1 μM, total protein but only fractionally active) (aa 106-492) proteolyzes peptidyl substrate boc-QAR-AMC (Shrimp, et al. 2020) (shown in red); however, following incubation with PAI-1 (1 nM active), soluble TMPRSS2 (aa 106-492) no longer exhibits enzymatic activity against the substrate (shown in blue). (B) Following incubation of PAI-1 with fractionally active, soluble TMPRSS2, a covalent complex between PAI-1 and TMPRSS2 is observed on a non-reducing western blot using a rabbit-anti‑PAI‑1 primary antibody.
