## Supplementary material for "High-throughput amino acid-level characterization of the interactions of plasminogen activator inhibitor-1 with variably divergent proteases": SI Fig. 2

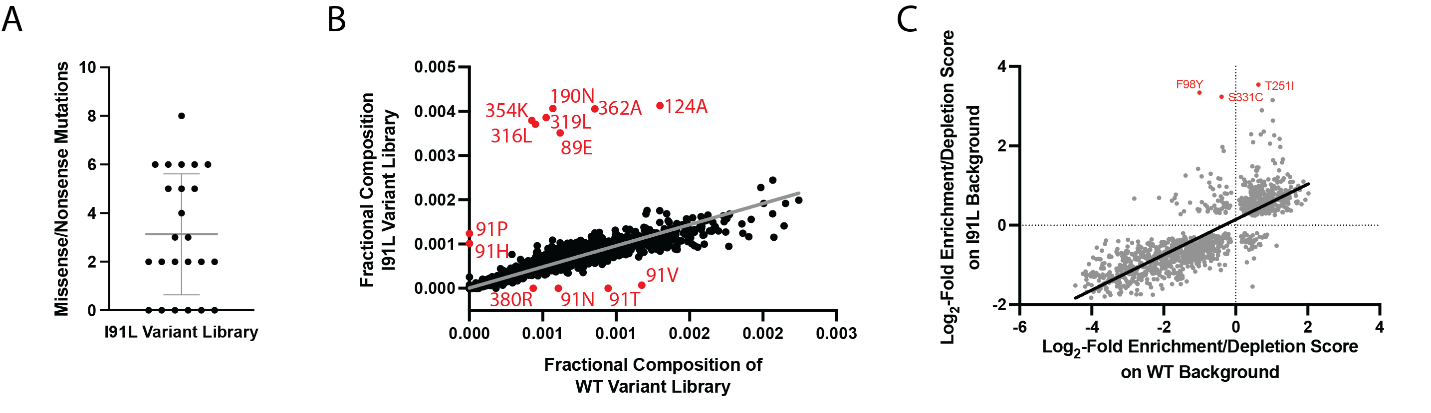

**SI Fig. 2. Validation of mutant library on the I91L PAI-1 backbone.** (A) The I91L PAI-1 variant library has a depth of 6.6 x 10^6^ unique clones with an average mutational frequency of 3.1 and a range of 0-8 missense/nonsense mutations per clone (individual points), as determined by Sanger sequencing of individual clones (n =24). (B) The fractional composition of our original WT PAI-1 variant library (Huttinger, et al. 2021; Haynes, et al. 2022) is compared to the I91L PAI-1 variant library used in the current study. The mutant composition of the libraries are highly correlated (slope = 0.95, R^2^ = 0.85, *grey line*). Amino acid substitutions that deviate from linearity are highlighted in *red*. Differences in the libraries at position 91 are likely due to codon usage and the different codon in the starting PAI-1 backbone (I or L at position 91). Other differences likely result from the random introduction of different mutations at early cycles of the error prone PCR. (C) Log_2_-fold enrichment/depletion scores for uPA inhibition by the PAI-1 variant library on the I91L background as a function of that on the WT PAI-1 background. The two libraries perform similarly to each other (slope = 0.45, R^2^ = 0.66, *black line*). PAI-1 variants further analyzed for their uPA and FXIIa inhibitory function (**Fig. 5**) are highlighted in *red*.
