## Supplementary material for "High-throughput amino acid-level characterization of the interactions of plasminogen activator inhibitor-1 with variably divergent proteases": SI Fig. 5

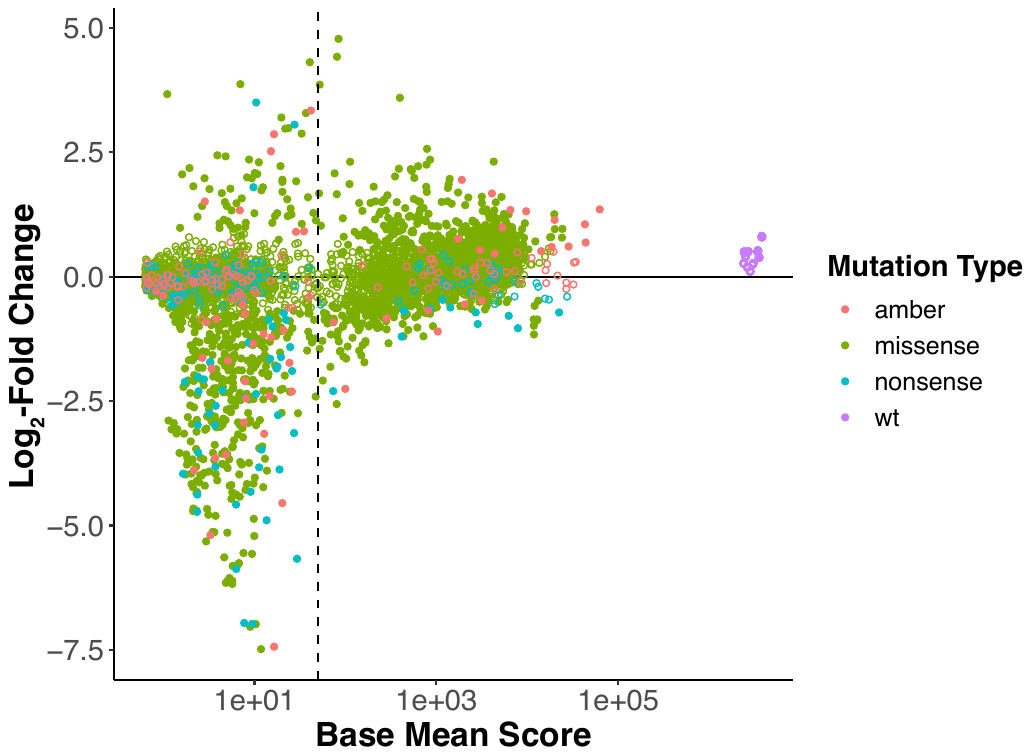

**SI Fig. 5. MA plot for DMS screen of PAI-1 inhibition of TMPRSS2.** The log_2_-fold change in the enrichment/depletion scores following the selection of the I91L PAI-1 variant library with TMPRSS2 is shown as a function of the BaseMean score (Love, et al. 2014). Variants with a p_adj_ > 0.1 are shown as solid circles, and variants with p_adj_ ≤ 0.1 are shown as open circles. Colors indicate the type of mutation: WT amino acids (*purple*), missense mutations (*green*), amber stop codon (*salmon*), and all other nonsense mutations (*blue*).
