## Supplementary material for "High-throughput amino acid-level characterization of the interactions of plasminogen activator inhibitor-1 with variably divergent proteases": SI Table 1

SI Table 1. Sequencing Primers

| Primer | 5’ → 3’ Sequence |
| --- | --- |
| pAY-FE PAI-1 Cloning Primers | |
| pAY-FE PAI-1 For | GCCGGCGCGCCAGAACAAAAACTCATCTCAGAAGAGGATC  TGGGTGGAGGTTCAGTGCACCATCCCCCATCCTAC |
| pAY-FE PAI-1 Rev | ACCTGCGGCCGCTGAACCACCTCCGGGTTCTATCACTTGGCCCAT |
| Sanger Sequencing/Colony PCR Primers | |
| pAYE Sequencing For | TTCATGCTGCCGGCTTTCTCGG |
| pAYE Sequencing Rev | TCGTCATCGTCCTTGTAGTCACCACCA |
| HTS Sequencing Primers | |
| PAI-1 Amplicon 1 For | NNNNNNGAACAAAAACTCATCTCAGAAGAGGATCTG |
| PAI-1 Amplicon 1 Rev | NNNNNNGGGTGAGAAAACCACGTTGCG |
| PAI-1 Amplicon 2 For | NNNNNNTTTCAGCAGGTGGCGCAG |
| PAI-1 Amplicon 2 Rev | NNNNNNCTTGTCATCAATCTTGAATCCCATAGCTGC |
| PAI-1 Amplicon 3 For | NNNNNNATGCTCCAGCTGACAACAGGA |
| PAI-1 Amplicon 3 Rev | NNNNNNTGTGGTGCTGATCTCATCCTTGTT |
| PAI-1 Amplicon 4 For | NNNNNNGCCCTCCGGCATCTGTACAAG |
| PAI-1 Amplicon 4 Rev | NNNNNNTTGCTTGACCGTGCTCCGG |
| PAI-1 Amplicon 5 For | NNNNNNAAGCTGGTCCAGGGCTTCATG |
| PAI-1 Amplicon 5 Rev | NNNNNNCCCAAGCAAGTTGCTGATCATACC |
| PAI-1 Amplicon 6 For | NNNNNNATTCATCATCAATGACTGGGTGAAGAC |
| PAI-1 Amplicon 6 Rev | NNNNNNCTGGAGTCGGGGAAGGGAG |
| PAI-1 Amplicon 7 For | NNNNNNAATGCCCTCTACTTCAACGGC |
| PAI-1 Amplicon 7 Rev | NNNNNNGGGCGTGGTGAACTCAGTATAG |
| PAI-1 Amplicon 8 For | NNNNNNCCCATGATGGCTCAGACCAAC |
| PAI-1 Amplicon 8 Rev | NNNNNNGGCAGAGAGAGGCACCTCT |
| PAI-1 Amplicon 9 For | NNNNNNAGCATGTTCATTGCTGCCCC |
| PAI-1 Amplicon 9 Rev | NNNNNNTTCAGTCTCCAGGGAGAACTTGG |
| PAI-1 Amplicon 10 For | NNNNNNCAGGCTGCCCCGCCT |
| PAI-1 Amplicon 10 Rev | NNNNNNACGTGGAGAGGCTCTTGGTC |
| PAI-1 Amplicon 11 For | NNNNNNAGACAGTTTCAGGCTGACTTCAC |
| PAI-1 Amplicon 11 Rev | NNNNNNGGGGGCCATGCGGGC |
| PAI-1 Amplicon 12 For | NNNNNNTCATCCACAGCTGTCATA |
| PAI-1 Amplicon 12 Rev | NNNNNNTGCGGCCGCTGAACCACCTCC |
